# Improved chromosome-level genome assembly for marigold (*Tagetes erecta*)

**DOI:** 10.1101/2023.07.25.550479

**Authors:** Fan Jiang, Lihua Yuan, Sen Wang, Hengchao Wang, Dong Xu, Anqi Wang, Wei Fan

**Affiliations:** Shenzhen Branch, Guangdong Laboratory for Lingnan Modern Agriculture; Genome Analysis Laboratory of the Ministry of Agriculture and Rural Affairs; Agricultural Genomics Institute at Shenzhen, Chinese Academy of Agricultural Sciences, Shenzhen, Guangdong, 518120, China; State Key Laboratory of Crop Stress Adaptation and Improvement, School of Life Sciences, Henan University, Kaifeng 475004, China; Shenzhen Research Institute of Henan university, Shenzhen 518000, China

**Author notes:** These authors contributed equally to this work (Fan Jiang, Lihua Yuan). Corresponding authors: Agricultural Genomics Institute at Shenzhen, Chinese Academy of Agricultural Sciences, Shenzhen 518120, China.

**Keywords:** *Tagetes erecta*, marigold, Asteraceae family, near telomere-to-telomere assembly, PacBio HiFi sequencing

## Abstract

Marigold (*Tagetes erecta* L.) is a popular ornamental plant of the Asteraceae family, and its petals are considered the most abundant source of lutein. A low-continuity chromosome-level genome sequence of marigold was published recently, with poor annotation of the protein-coding genes, which hinders the studies of lutein biosynthesis. Here, we generated a near telomere-to-telomere level genome assembly of marigold based on highly accurate high-fidelity (HiFi) long reads and Hi-C sequencing data. Compared to the previously reported marigold genome, the current assembly had obviously higher contiguity and higher completeness of gene set. The current genome assembly has a 27-fold increase in contig N50 size, a 12.1% increase in chromosome anchoring rate, and a 9.0% increase in BUSCO complete rate for the gene set. Besides, the current assembly has much fewer assembly errors. Based on this high-quality genome assembly, we found that the 170-bp repeats are the most abundant centromeric unit and all centromeric regions are distributed along the whole chromosomes for all 12 centromeres, indicating the existence of the holocentromeres in marigold. In addition, we analyzed the structure and phylogenetic relationship of the four *PSY* genes, and revealed that these genes have diversified and possibly executed different functions in various tissues. Our near telomere-to-telomere level genome assembly and comprehensive gene annotation will greatly facilitate the breeding of marigold and researches aimed at improving lutein production.

## 1. Introduction

Marigold (*Tagetes erecta* L.) is one of the most popular ornamental plants of the Asteraceae family, and its flower petals has become the most important raw material for lutein production (Lin et al., 2015; Zhang et al., 2020b; Zhang et al., 2020a). Marigold is an annual herbaceous plant with yellow or orange flowers (Burlec et al., 2021; Moliner et al., 2018), and its flower petals are considered the most abundant source of lutein (Nwachukwu et al., 2016; Park et al., 2017; Tanaka et al., 2008), which is effective in the prevention and treatment of cancer, coronary artery disease, and many eye diseases (Buscemi et al., 2018; Chung et al., 2017; Gansukh et al., 2019; Li et al., 2020).

The lutein biosynthetic pathway has been well studied in many high plants, and it starts with the condensation of two geranylgeranyl diphosphate (GGPP) molecules to produce phytoene, which is catalyzed by phytoene synthase (PSY) (Nisar et al., 2015; Sun et al., 2018). Subsequently, phytoene is transformed to lycopene by a series of desaturation and isomerization reactions (Giuliano, 2014). Then, lycopene is cyclized twice by lycopene *ε*-cyclase and lycopene *β*-cyclase to generate *α*-carotene, which is hydroxylated to lutein by hydroxylase (Kim et al., 2009). In this pathway, PSY catalyzes the first and rate-limiting step in lutein biosynthesis (Maass et al., 2009; Nisar et al., 2015; Rodriguez-Villalon et al., 2009), and four *PSY* genes in marigold were identified on the basis of a whole genome sequence (Xin et al., 2023). However, the functional differences of these genes among roots, stems, leaves, and flowers in marigold was less understood (Feng et al., 2018).

A recently study by Xin et al has reported a low-continuity marigold genome assembly, which contains 3,451 contigs with a contig N50 size of 1.4 Mb, and only 87.6% of contig sequences were anchored to 12 pseudo-chromosomes (Xin et al., 2023). In addition, the ratio of transposable elements (57.3%) and BUSCO complete rate (89.3%) for the genes were obviously lower than those of the closely related species (Badouin et al., 2017; Fan et al., 2022; Liu et al., 2020; Xu et al., 2021). In this study, we generated a near telomere-to-telomere level reference genome assembly for marigold, explored the structure of centromeres, and investigated the expression patterns of the *PSY* genes in different tissues.

## 2. Materials and Methods

### 2.1. Plant materials and genome sequencing

The seeds of marigold were provided by Suqian Sunrise Seed Industry Co., Ltd, in March 2022, and then were grown in a greenhouse of Agricultural Genomics Institute at Shenzhen, Chinese Academy of Agricultural Sciences, Guangdong province, China. Total genomic DNA was extracted from fresh leaves using the Hi-DNAsecure Plant Kit (TIANGEN, China). Then, sequencing libraries were constructed by SMRTbell Express Template Prep Kit 2.0 (PacBio, USA), and sequenced on a PacBio Sequel II platform with Circular consensus sequence (CCS) mode (PacBio, USA). For Hi-C sequencing, the cross-linked DNA sample was digested with the MboI at motifs GATC, and paired-end sequenced (2 x 150 bp) on an Illumina NovaSeq 6000 platform.

### 2.2. Transcriptome sequencing

To assist the prediction of protein-coding genes, total RNA samples were extracted by RNeasy Plant Mini Kit (QIAGEN, Germany) from the different tissues, and then were reverse-transcribed into cDNA using NEBNext Single Cell/Low Input cDNA Synthesis & Amplification Module (NEB, UK). Sequencing libraries were constructed by SMRTbell Express Template Prep Kit 2.0 (PacBio, USA) and sequenced on PacBio Sequel II system (PacBio, USA).

To investigate the expression patterns of candidate genes involved in lutein biosynthesis, the samples collected from the roots, stems, leaves and flowers of 80-day-old plants were used for tissue specificity analysis. Each plant represents a biological repeat, and five biological replicates were used in this study. All samples were immediately frozen in liquid nitrogen and stored at -80°C until use. Total RNA for each sample was extracted using RNeasy Plant Mini Kit (QIAGEN, Germany) according to the manufacturer’s instructions, and the cDNA libraries were paired-end (2 x 150 bp) sequenced on an Illumina NovaSeq 6000 system.

### 2.3 Genome assembly

Before genome assembly, the PacBio HiFi reads with read length < 2,000 bp or average quality < 99% were filtered out. Then, the genome size was estimated by GCE v1.02 (Liu et al., 2013) based on k-mer frequencies (K = 17). The filtered PacBio HiFi reads were assembled using Hifiasm v0.16.1 (Cheng et al., 2021) with parameter “-l 3”. Purge_dups v1.2.5 (Guan et al., 2020) with parameters “-a 83 -f 0.8 -e” was used to filter the duplicate contigs in the primary assembly. After removing contaminant contigs, we obtained 47 contigs with a total length of 778.9 Mb and a contig N50 size of 38.0 Mb (**Table 1**). The completeness of the assembled contigs were evaluated using the BUSCO v5.2.2 (Simao et al., 2015) with embryophyta_odb10 database, and the contig correctness was estimated based on yak QV (https://github.com/lh3/yak).

**Table 1.**
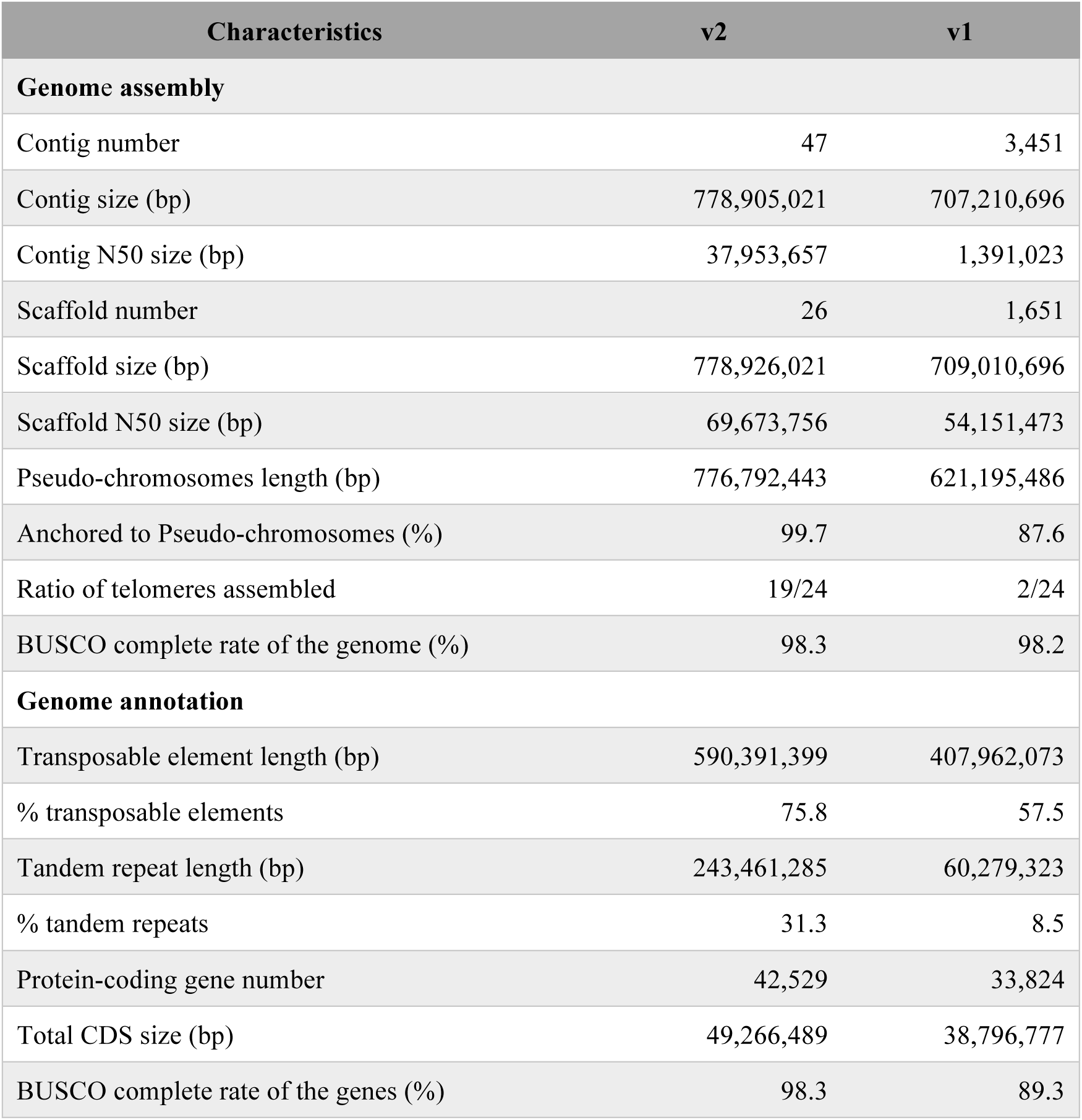
Comparisons of the genomic assembly and annotation for marigold v2 and v1 assembly.

The pseudo-chromosomes were constructed using Hi-C reads. We first mapped the Hi-C reads onto the assembled contigs by Bowtie2 v2.3.4.3 (Langmead and Salzberg, 2012), and then used HiC-Pro v2.11.4 (Servant et al., 2015) pipeline to detect valid ligation pairs and generate the Hi-C link matrixes among different contigs. At last, the contigs were clustered, ordered, and oriented into pseudo-chromosomes using EndHiC v1.0 (Wang et al., 2022) with parameters “--rounds 3 -- binnumstep 6”.

### 2.4. Transposable elements identification

Transposable elements (TEs) were identified through three steps: (1) we used EDTA v1.9.9 (Ou et al., 2019) to produce a filtered TE library; (2), we performed homology-searching against the above structural TE library, Repbase v26.05 (Bao et al., 2015), and protein-coding TE database using RepeatMasker v4.1.2 (Smit et al.); (3) we masked the above identified TEs with Ns in the genome, and used RepeatModeler v2.0.2 (Flynn et al., 2020) to construct an extra *de novo* TE library, which was further classified by TERL (da Cruz et al., 2021). All classified TE sequences in the *de novo* TE library were used by RepeatMasker to identify the remaining TEs in the genome.

### 2.5. Gene prediction and functional annotation

We used Augustus v3.4.0 (Stanke et al., 2006) to predict the protein-coding gene models. The training parameters for Augustus were generated during the BUSCO (Simao et al., 2015) completeness assessment of the genome assembly. The homology hints were generated by aligning the protein-coding sequences from the published genomes of *Smallanthus sonchifolius* (Fan et al., 2022), *Mikania micrantha* (Liu et al., 2020), *Helianthus annuus* (Badouin et al., 2017), and *Stevia rebaudiana* (Xu et al., 2021) to marigold genome, using Exonerate v2.2.0 (Slater and Birney, 2005). To produce transcription hints, the full-length transcripts and Illumina RNA-Seq reads were aligned to the genome, and then were transformed to hints by blat2hints and bam2hints in Augustus, respectively. The tRNA genes were predicted by tRNAscan-SE v2.0 (Chan et al., 2021), and the rRNA genes were predicted by cmscan from infernal v1.1.4 (Nawrocki and Eddy, 2013). Functional annotation was performed by searching the protein sequences against the NCBI-NR and KEGG databases using DIAMOND v2 (Buchfink et al., 2021). The protein domain annotation was performed using InterProScan v5.52-86 (Blum et al., 2021) against InterPro database.

### 2.6. Identification of telomeres and centromeres

To investigate the telomeres and centromeres, tandem repeats were explored using Tandem Repeats Finder (TRF) v4.07 (Benson, 1999) with parameter “2 5 7 80 10 50 2000 -h -d”. We identified telomeres on the chromosome scaffolds based on TRF results and telomeric units of “TTTAGGG” and “CCCTAAA”. For centromere annotation, we investigated the tandem repeat units with length of 100 - 200 bp, and counted the total length of each repeat units, revealing the 170-bp repeats were the most abundant unit in the genome, followed by 167-bp, 178-bp and 169-bp repeats. The distribution of the four repeat units in different pseudo-chromosomes were visualized by IGV v2.11.4 (Thorvaldsdottir et al., 2013).

### 2.7. Analysis of genes involved in lutein biosynthesis

The lutein biosynthesis genes in marigold genome were identified by homology alignment. Firstly, we downloaded the known genes involved in the lutein biosynthesis pathway (map00906) from KEGG database for the closely related species in Asterales order, including *Helianthus annuus*, *Erigeron canadensis*, *Lactuca sativa*, and *Cynara cardunculus* var. *scolymus*. Then, we aligned the protein-coding genes of marigold to the downloaded lutein biosynthesis genes using DIAMOND v2 (Buchfink et al., 2021) with parameters “blastp --more-sensitive --evalue 0.00001”. The genes in marigold with best alignment identity > 80% and coverage > 80% were taken as potential lutein synthesis genes. Furthermore, the orthogroups constructed by OrthoFinder v2.5.2 (Emms and Kelly, 2019) that contained these genes were obtained and checked.

To construct the tree for *PSY* genes, we downloaded the protein sequences of *PSY* genes from the KEGG database for the 4 Asterales species (*Helianthus annuus*, *Erigeron canadensis*, *Lactuca sativa*, *Cynara cardunculus* var. *scolymus*, and *Solanum lycopersicum*) and 4 Solanales species (*Solanum lycopersicum*, *Solanum pennellii*, *Solanum stenotomum*, and *Ipomoea triloba*). The tree was constructed by fasttree v2.1.11 (Price et al., 2010) with parameters “-lg -gamma” based on multiple sequences alignments from protein sequences. The expression level of these candidate genes was evaluated by transcripts per million (TPM) calculated from the HISAT v2.2.1 (Kim et al., 2019) mapping results of root, stem, leaf, and flower transcriptome data.

## 3. Results

### 3.1. Near telomere-to-telomere level genome assembly of marigold

A total of 49.7 Gb (64 ×) PacBio HiFi reads with a read N50 length of 18.3 Kb (**Supplementary Table S1**) were assembled into contigs using Hifiasm (Cheng et al., 2021). The primary assembly includes 381 contigs with a total length of 1,011 Mb, a contig N50 size of 36.5 Mb, and a contig N90 size of 8.8 Mb (**Supplementary Table S2**). This assembly is obviously larger than the estimated genome size (∼780 Mb) that evaluated based on the distribution of k-mer frequencies (**Supplementary Figure S1**), which may result from genome heterozygosity. After purging the heterozygous contigs from the primary assembly using Purge_dups (Guan et al., 2020), we obtained a genome assembly with a total length of 778.9 Mb (**Table 1**), which is comparable to the estimated genome size. The genome assembly comprises 47 contigs with contig N50 and N90 size of 38.0 and 11.0 Mb, respectively (**Table 1 and Supplementary Table S2**), revealing a BUSCO complete rate of 98.3% (**Table 1**). In addition, the contig correctness was evaluated by using yak, which revealed an average base accuracy of 99.9998% (QV57), suggesting the high accuracy of our genome assembly. Compared to recently published genome assembly of Marigold by Xin et al. (defined as assembly v1) (Xin et al., 2023), the huge improvement of our genome assembly (defined as assembly v2) was highlighted by the total contig number (∼73-fold decrease), contig N50 size (27-folds increase), and genome size (71.7 Mb increase) (**Table 1**). Based on the Hi-C data (**Supplementary Table S1 and S3**), we successfully anchored 99.7% (776.8 Mb) of contig sequences onto 12 pseudo-chromosomes (**Table 1**, **Figure 1a and b**), which was obviously higher than that of v1 assembly (87.6%) (**Table 1**). Finally, we obtained a near telomere-to-telomere level reference genome for marigold, with scaffold N50 size of 69.8 Mb and N90 size of 40.8 Mb.

**Figure 1.**
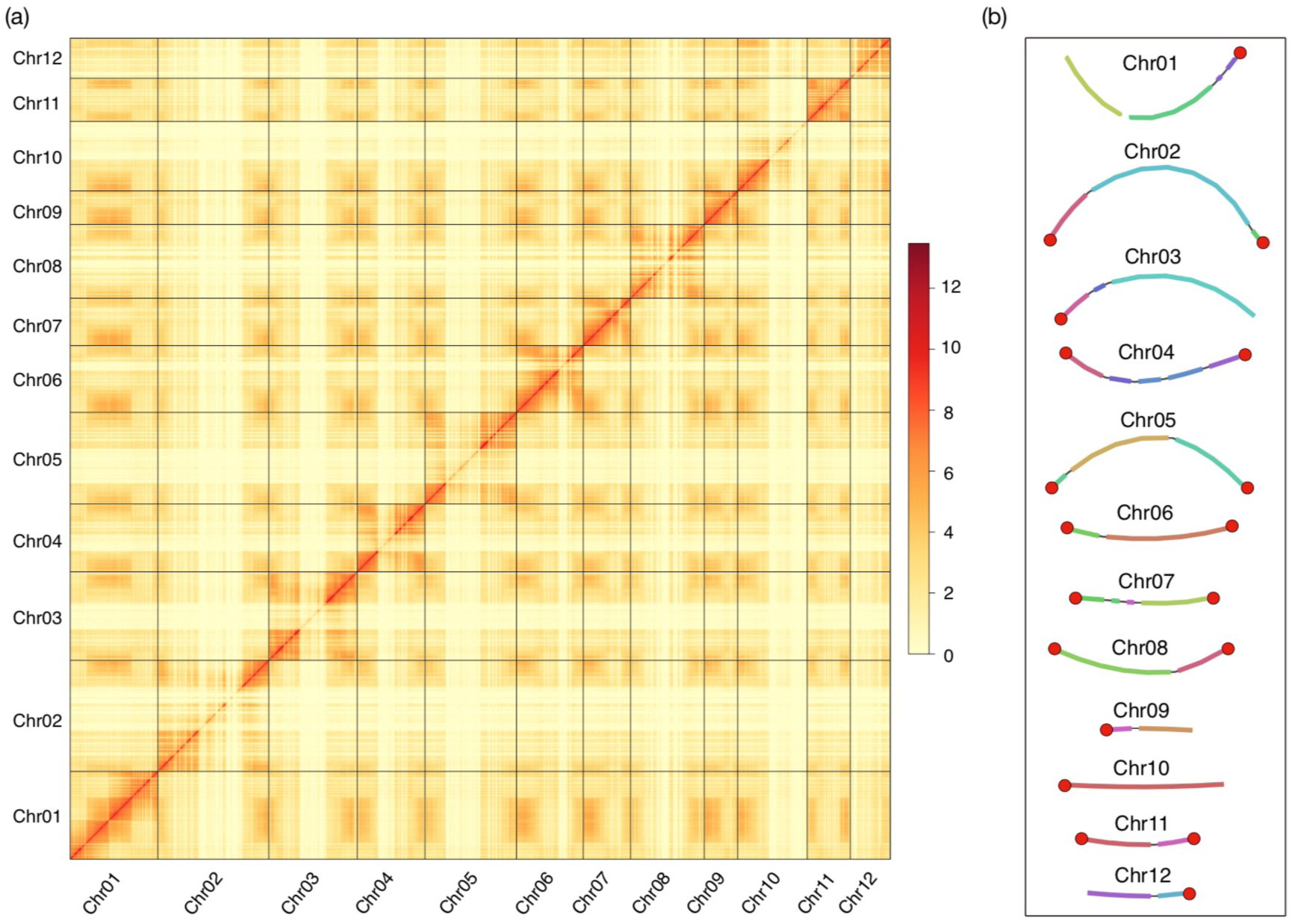
Assessment of the genome assembly for marigold. (a) Hi-C heatmap of the genome assembly. We scanned the genome by 200-Kb non-overlapping window as a bin and calculated valid interaction links of Hi-C data between any pair of bins, and color represents Log2(links number); (b) View of the pseudo-chromosomes. The thick lines represent contigs, and the thin lines represent the links between the two contigs. The chromosome ends assembled with telomere-specific repeats (CCCTAAA) were highlighted with red solid circle.

We identified a total of 590.4 Mb (75.8%) of TEs in v2 assembly (**Table 1 and Supplementary Figure S2**), including 84.1 Mb (14.2%) of structural intact TEs. The proportion of TEs in v2 assembly falls in the range of that for the closely related species (73.4% - 84.7%) (Badouin et al., 2017; Fan et al., 2022; Liu et al., 2020; Xu et al., 2021), and showed a 18.1% increase compared to v1 assembly (57.7%) (**Table 1 and Supplementary Table S4**). In addition, we identified a total of 242.8 Mb (31.2% of the genome) TRs in marigold (**Table 1**). Then, we compared the TEs and TRs in v2 assembly, and found that 97.5% of TRs were overlapped with TEs (**Supplementary Figure S3**), which means that most of the TRs were formed from tandem clusters of the same type of TEs.

We predicted 42,529 protein-coding genes in v2 assembly, similar to that of *Mikania micrantha* (46,351), *Helianthus annuus* (57,126), and *Stevia rebaudiana* (44,143), and obviously fewer than that of *Smallanthus sonchifolius* (89,959) (Badouin et al., 2017; Fan et al., 2022; Liu et al., 2020; Xu et al., 2021), which experienced a recent whole genome duplication (Fan et al., 2022). The genome has a mean coding sequence length of 1,158 bp and exon number of 5.1, which was similar to that of the related species (**Supplementary Table S5**). Functional annotation showed that 86.7% of genes could be annotated by one of the NCBI-NR, KEGG, InterPro or GO databases (**Supplementary Table S6**). Compared to v1 assembly, v2 assembly identified more genes (8,705 genes), which resulted in a 9.0% increase in BUSCO complete rate (**Table 1 and Supplementary Table S5**), indicating the higher completeness of the v2 gene set. In addition, we identified 1,098 tRNA genes, 2,126 rRNA genes, and 929 other non-coding RNA genes in v2 assembly (**Supplementary Table S7**).

### 3.2. Comprehensive assembly of repetitive regions and correction of large-scale assembly errors

In v2 assembly, 79.2% of the assembled chromosome-ends have telomeric repeats (**Table 1**), including 7 pseudo-chromosomes have telomeric repeats at both ends and 5 pseudo-chromosomes have telomeric repeats at a single end (**Figure 1b and Supplementary Figure S4**). Compared to v1 assembly, the v2 assembly reconstructed obviously more telomeres (79.2% vs 8.3%) (**Table 1**), indicating the improvement in assembly of the ends for pseudo-chromosomes.

Compared to v1 assembly, the v2 assembly has an obvious increase (155.6 Mb) in the total length of the 12 pseudo-chromosomes, which caused by both the higher total contig size (778.9 Mb vs 707.2 Mb) and the higher anchor rate (99.7% vs 87.6%) in v2 assembly (**Table 1**). Comparisons of the pseudo-chromosomes between the assembly v2 and v1 revealed an obviously one-to-one match relationship for all 12 pseudo-chromosomes (**Figure 2**). Compared to v2 assembly, large blocks were missed in v1 assembly for many pseudo-chromosomes, including Chr02 (33.1 Mb), Chr03 (27.5 Mb), Chr04 (9.1 Mb), Chr05 (32.4 Mb), Chr06 (10.0 Mb), Chr08 (20.1 Mb), and Chr10 (25.5 Mb) (**Figure 2 and Supplementary Table S8**). After annotating of these large blocks in v2 assembly, we found that most of these sequences were identified as TRs, which were barely assembled in v1 assembly (**Figure 2**).

**Figure 2.**
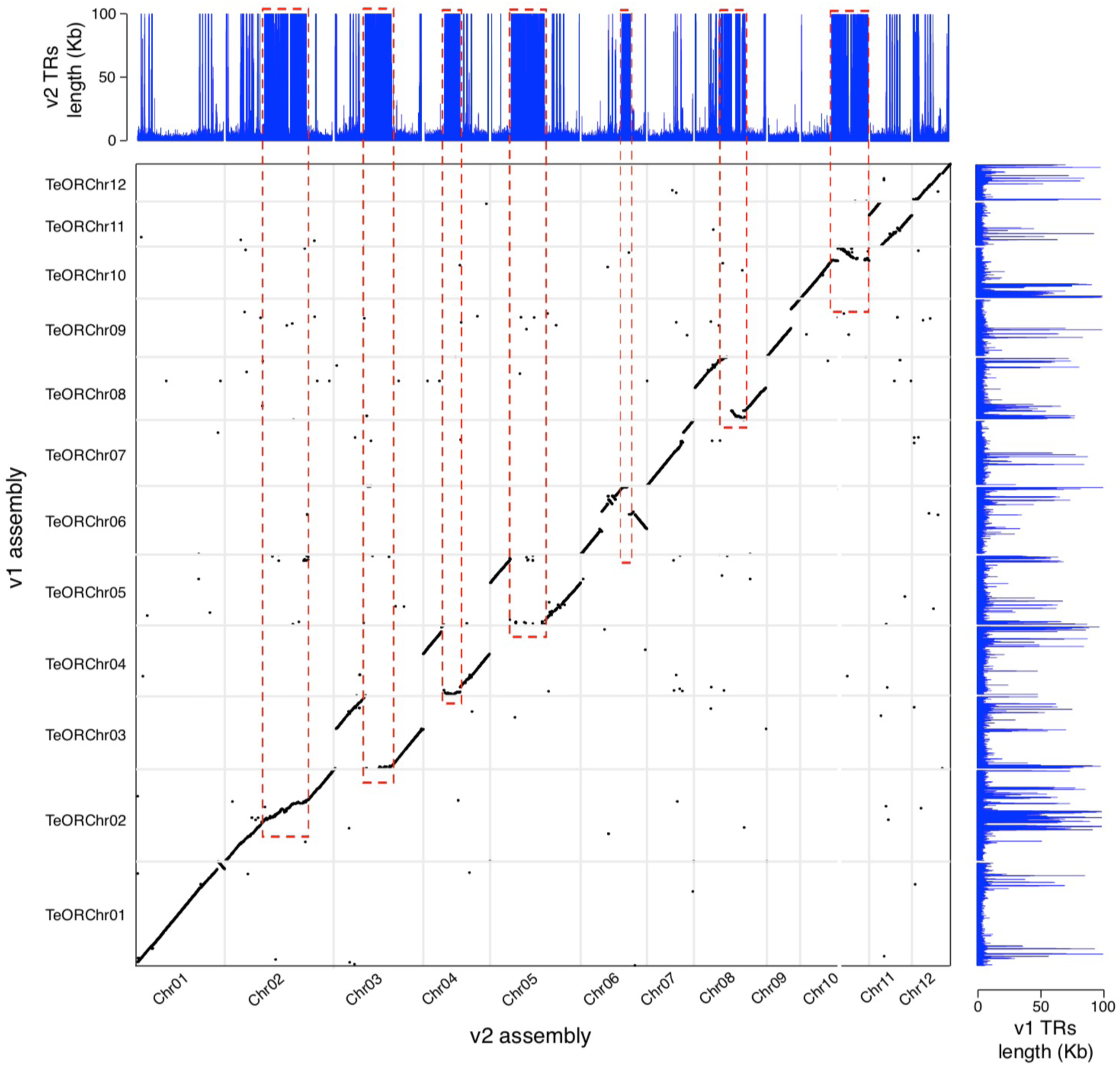
Comparisons between the genomes of marigold v2 and v1 assembly. The above and right parts showed the distributions of tandem repeats in v2 and v1 assembly, respectively. We scanned the genome by 100-Kb non-overlapping window as a bin to calculate the length of tandem repeats. The core part of the figure showed the pair-wise alignment of genome sequences between the v2 and v1 assembly. The corresponding segments for highly enriched tandem repeats in v2 assembly were highlighted with red box.

Moreover, the pseudo-chromosomes with large blocks that showed inconsistent ordering or orientation in v1 assembly were analyzed. For Chr03, the whole pseudo-chromosome in v1 assembly was split into two large blocks by a huge gap according to the alignment result, and the ordering of the two blocks was inconsistent with that in v2 assembly (**Figure 2**). Similar results were observed for Chr04, Chr05, and Chr08 (**Figure 2**). Because the huge gap will greatly reduce the Hi-C linkage signal between the two blocks (Zhang et al., 2019), it is easy to result in wrong connections among different chromosome-ends in v1 assembly. In addition, no telomeric repeats was identified from the ends of these pseudo-chromosomes in v1 assembly, but identified from the single or both ends in v2 assembly (**Figure 1b and Supplementary Figure S4**), which suggested the correctness of these pseudo-chromosomes in v2 assembly. For Chr01, Chr06 and Chr11, the chromosomes in v1 assembly were split into many large blocks, which were inconsistent with that in v2 assembly, though the length of these chromosomes are similar (**Figure 2**). Since v1 assembly has telomeric repeats located in the sub-terminal of the three pseudo-chromosomes instead of the chromosome-ends, while v2 assembly has telomeric repeats at single or both ends (**Figure 1b and Supplementary Figure S4**), we inferred that the v2 assembly has corrected the order and orientation mistakes in v1 assembly. In total, the v2 assembly corrected 7 pseudo-chromosomes that contained wrong ordering or orientation blocks in v1 assembly.

### 3.3. High proportion of centromeric 170-bp unit reveals the existence of holocentromeres

The Hi-C heatmap showed many large blocks in genome assembly with no or weak Hi-C linkage signals (**Figure 1a**), revealing a high proportion of high similar repeating sequences existed in the genome. We identified a total of 242.8 Mb (31.2% of the genome) TRs, and the distribution of the TRs was highly consistent with the location of these large blocks in Hi-C heatmap (**Supplementary Figure S5**). Noticeably, the proportion of TRs in marigold genome was obviously higher than that of *Smallanthus sonchifolius* (101.3 Mb, 3.7% of the genome), *Mikania micrantha* (108.6 Mb, 6.1% of the genome), *Helianthus annuus* (163.1 Mb, 5.4% of the genome), and *Stevia rebaudiana* (72.8 Mb, 5.1% of the genome). In addition, the TRs and TEs were highly overlapped in marigold genome (**Supplementary Figure S3**), and more than 97.5% of TRs were classified as TEs.

Previous studies have reported that centromere consisted of highly repetitive DNA sequences with monomers of length 100 - 200 bp (Melters et al., 2013; Naish et al., 2021; Shi et al., 2023; Sundararajan and Straight, 2022). Here, we found that 170-bp repeats were the most abundant unit in the genome, which had 682,138 repetitions accounted for 14.8% of the total genome sequences, followed by 167-bp (2.7%), 178-bp (1.2%) and 169-bp (0.9%) repeats (**Supplementary Table S9**). These four repeat units localized on all 12 pseudo-chromosomes, with the total length ranged from 4.0 Kb (Chr09) to 29.0 Mb (Chr02) (**Supplementary Table S10**). In detail, the 170-bp was the main repeat unit in Chr01, Chr02, Chr03, Chr05, Chr06, and Chr07, and the 169-bp was the main repeat unit in Chr11 (**Figure 3a and Supplementary Figure S6**). In addition, the 167-bp and 170-bp repeats were the main units for Chr04, Chr08, and Chr10, the 167-bp and 178-bp repeats were the main units for Chr09, and 169-bp and 170-bp repeats were the main units for Chr12 (**Figure 3a and Supplementary Figure S6**). Noticeably, these repeat units distributed across the whole chromosome for all 12 pseudo-chromosomes, but not constrained in a single region (**Figure 3b**). Besides, these repeat regions have obviously lower gene density than that of other parts on all pseudo-chromosomes (**Figure 3b**). Moreover, more than 80.2% of sequences within these repeat regions were annotated as retrotransposons (**Figure 3b**), which was consistent with that centromeres are composed of long arrays of tandem repeats and/or retrotransposons in many species (Henikoff et al., 2001; Zhang et al., 2014). Based on these results, we inferred that these repeat regions in marigold genome were centromeric regions, and the chromosomes of marigold should be classified as holocentric chromosomes, similar to *Holhymenia histrio*, *Rhynchospora pubera*, and *Vitis vinifera* (Bardella et al., 2020; Hofstatter et al., 2022; Marques et al., 2015; Shi et al., 2023).

**Figure 3.**
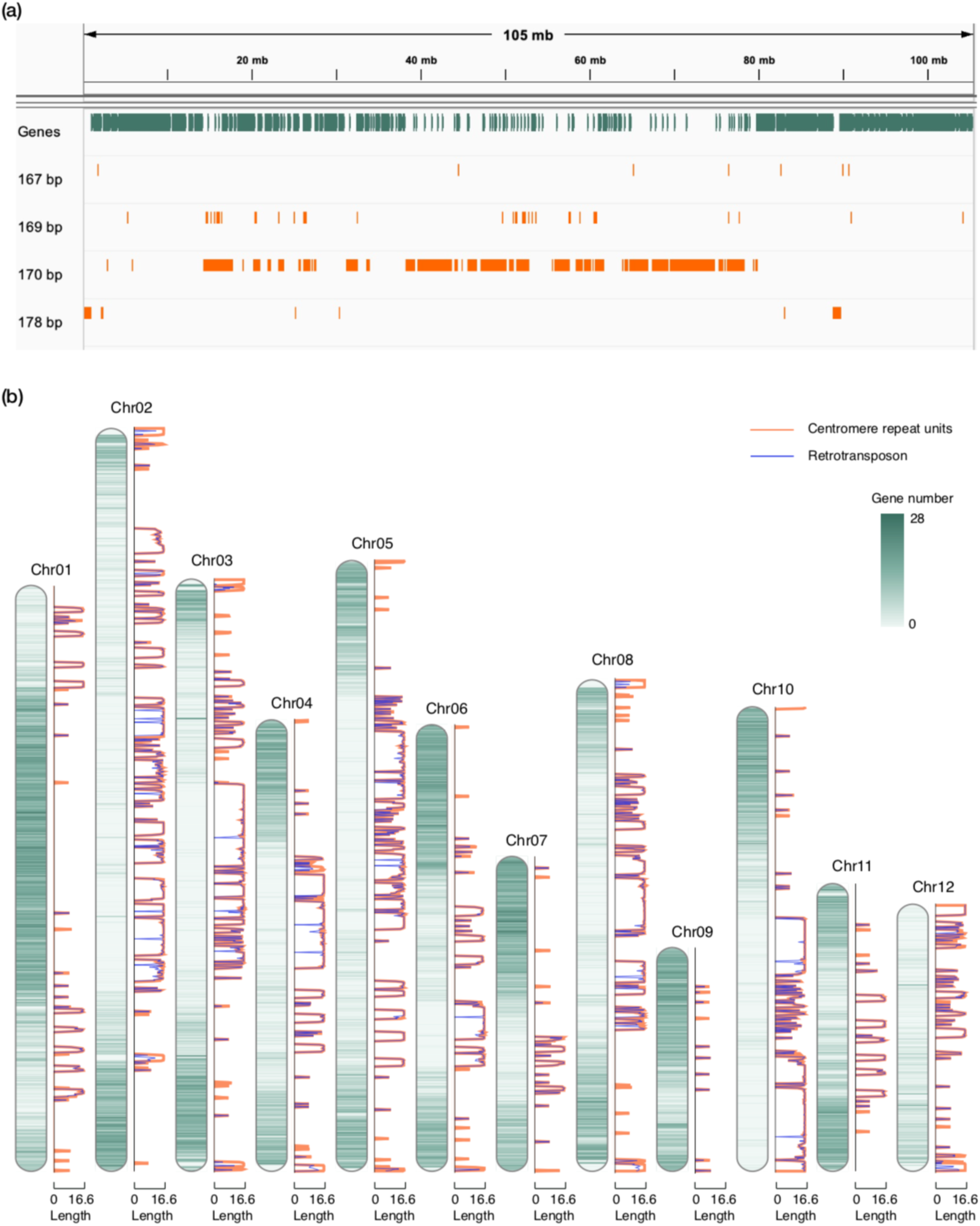
Characterization of holocentromeres in marigold genome. (a) Views of genes and four centromeric repeat units for Chr02 with IGV. We take Chr02 as an example to show the distributions of genes and different centromeric units. The head number showed the total length of the chromosome. (b) Distribution of genes, centromeric repeat units and the retrotransposons in centromeric regions for all 12 pseudo-chromosomes. The left columns showed the distribution of genes. The length represents Log2(length) in 100 Kb windows.

### 3.4. Differential expression patterns of the four *PSY* genes in different tissues of marigold

In marigold, lutein accounts for up to 90% of the total carotenoid contents in the flower petals, and all genes involved in lutein biosynthesis were identified by Xin *et al*. based on a whole genome sequence (Xin et al., 2023). Phytoene synthase (PSY) catalyzes the first and rate-limiting step in lutein biosynthesis (**Figure 4a**), and four *PSY* genes were identified from marigold genome (Xin et al., 2023). However, the structure and the phylogenetic relationship of the four genes in marigold were not investigated. Based on our high-quality genome assembly, we identified four *PSY* genes (*PSYa*, *PSYb*, *PSYc*, and *PSYd*) (**Supplementary Figure S7**), consistent with the previous study (Xin et al., 2023). The four *PSY* genes have a length of 2,841 bp, 2,760bp, 3,235 bp, and 1,917 bp, respectively, all of which contain 6 exons and 5 introns (**Supplementary Figure S7**). The predicted protein sequences have 425, 398, 427, and 375 amino acids for *PSYa*, *PSYb*, *PSYc*, and *PSYd* genes, respectively. To investigate the phylogenetic relationship of these four *PSY* genes, 28 PSY protein sequences from four Solanales species and five Asterales species were used (**Figure 4b**). The phylogenetic tree showed that *PSYa* and *PSYc* were tightly grouped together, which was then clustered with *PSYb* (**Figure 4b**). In addition, *PSYd* was clustered into another group, and showed a high divergence to other three *PSY* genes (**Figure 4b**), which was consistent with that *PSYd* showed a low amino acid sequence identity to *PSYa*, *PSYb* and *PSYc* (60.9 - 62.2%).

**Figure 4.**
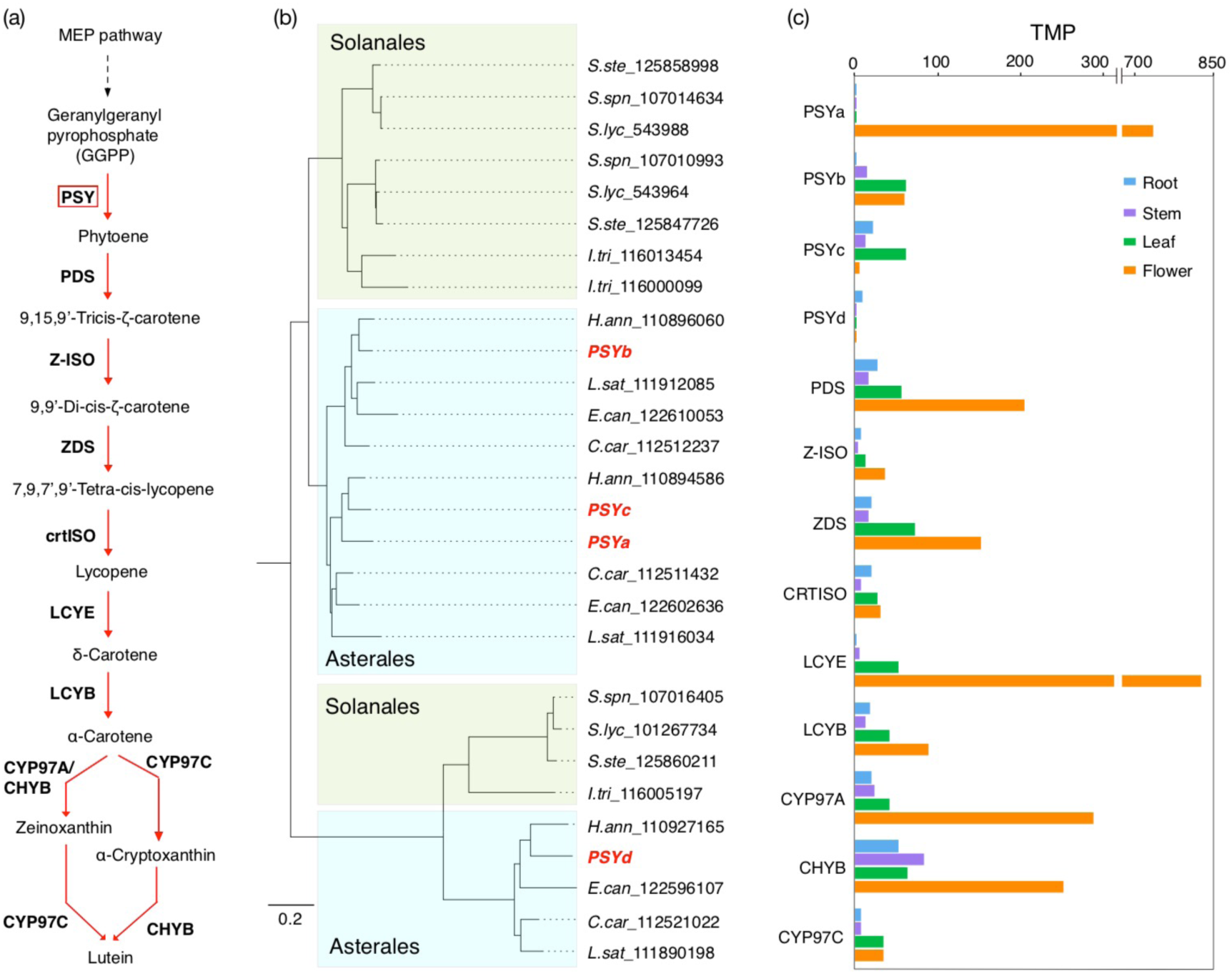
Analysis of lutein biosynthesis in marigold. (a) Primary steps for lutein biosynthesis pathway. (b) Phylogenetic tree for *PSY* genes from some Solanales and Asterales species. (c) Comparisons of the expression levels of all genes involved in lutein biosynthesis among different tissues.

Based on the recently published marigold genome, Xin *et al*. investigated the expression patterns of lutein biosynthesis genes at four flowering development stages in marigold flowers (Xin et al., 2023), but the differences among different tissues were not studied. In this study, we compared the expression levels of these genes among different tissues, including roots, stems, leaves, and flowers. For *PSY* genes, *PSYa* only expressed in flowers and showed a high expression level, *PSYb* showed moderate expression levels in flowers and leaves, and low level in stems, and *PSYc* exhibited moderate expression levels in leaves, and low level in stems and roots (**Figure 4c**). In addition, *PSYd* only expressed in roots with very low expression level (**Figure 4c**), which was similar to its homolog *SlPSY3* in tomato that showed low but detectable transcript levels in roots (Fantini et al., 2013). These results suggested that the four *PSY* genes have diversified and possibly executed different functions in roots, stems, leaves, and flowers. In addition, the expression levels of the other genes were all obviously higher in flowers than those in leaves, stems, and roots (**Figure 4c**), which was consistent with that the flowers accumulated with the highest level of lutein in marigold (Moehs et al., 2001).

## 4. Discussions

In this study, we reported a near telomere-to-telomere level genome assembly of marigold based on highly accurate HiFi long reads and Hi-C sequencing data. Compared to the previously reported marigold genome, the current assembly had obviously higher contiguity and higher completeness of gene set. Besides, the current assembly has much fewer assembly errors. Although all the assembled pseudo-chromosomes contain less than 5 contigs in this study, a few gaps still exist in the genome assembly. In addition, we have assembled telomere sequences for 19 chromosome-ends, but 5 chromosome-ends still have no assembled telomere sequences. To generate a real telomere-to-telomere genome assembly, further investigations based on ultra-long sequencing and other technologies will be necessary.

Based on the high-quality genome assembly, we found that the 170-bp repeats are the most abundant centromeric unit, which followed by 167-bp, 178-bp and 169-bp repeats. All centromeric units are distributed along the whole chromosomes for all 12 centromeres, indicating the existence of the holocentromeres in marigold. Here, the centromeres were identified mainly based on the analysis of the tandem repeat units. To ascertain the centromere type of marigold, more studies are needed to investigate the organization of spindle fibres at metaphase, since the previous studies reported that the spindle fibres are attached along almost the entire poleward surface of the chromatids in holocentromeres (Schubert et al., 2020). In addition, the chromosomal distribution of the centromere-specific histone H3 (CENH3) protein and the cell cycle-dependent pericentromeric phosphorylation of histone H3 and H2A need further investigations (Baez et al., 2020).

The reference genome sequence of marigold greatly facilitates lutein biosynthesis studies. Xin *et al*. have identified all the genes involved in lutein biosynthesis based on a low-continuity reference genome of marigold, and investigated the expression patterns of these genes at four flowering development stages in flowers (Xin et al., 2023). However, in marigold, the relationship of the four *PSY* genes, and the expression patterns of these genes among different tissues were not studied. In this study, we analyzed the structure and phylogenetic relationship of the four *PSY* genes, revealing that *PSYa* and *PSYc* were tightly grouped together, then clustered with *PSYb*, and *PSYd* formed a separate group that with a high divergence to other three *PSY* genes. In addition, our results revealed that the four *PSY* genes have diversified and possibly executed different functions in roots, stems, leaves, and flowers. To verify the functions of these *PSY* genes, more investigations based on the molecular genetic methods, such as gene editing technologies, are needed. Our near telomere-to-telomere level genome assembly and comprehensive gene annotation will greatly facilitate the breeding of marigold and researches aimed at improving lutein production.

## Data Availability Statement

All raw sequencing data generated during the current study have been deposited at DDBJ/ENA/GenBank under project accession PRJNA975275. Genomic sequence reads have been deposited in the SRA database with accession SRR24689467 for PacBio sequencing. Hi-C sequencing reads have been deposited in the SRA database with accession SRR24718810. Full-length transcript sequence reads have been deposited in the SRA database with accession SRR24733750. Illumina transcript sequences for roots, stems, leaves, and flowers have been deposited in SRA database with accession SRR24733730-SRR24733749. The genome assembly and annotation have deposited at NCBI with accession JAUHHV000000000, and also available at China National Genomics Data Center (https://ngdc.cncb.ac.cn) with the Project ID PRJCA017901 and Zenodo: https://www.zenodo.org/record/8107331.

## CRediT authorship contribution statement

**Fan Jiang**: Investigation, Methodology, Formal analysis, Writing - original draft. **Lihua Yuan**: Investigation, Methodology, Formal analysis. **Sen Wang**: Formal analysis, Validation. **Hengchao Wang**: Formal analysis. **Dong Xu**: Formal analysis. **Anqi Wang**: Validation, Software. **Wei Fan**: Supervision, Writing - review & editing, Funding acquisition.

## Acknowledgment

The work was funded by the Agricultural Science and Technology Innovation Program and the Elite Young Scientists Program of CAAS, the fund of Key Laboratory of Shenzhen (ZDSYS20141118170111640), and the National Natural Science Foundation of China (32172430).

