## Supplementary_files for "Improved chromosome-level genome assembly for marigold (*Tagetes erecta*)"

**Supplementary figures**

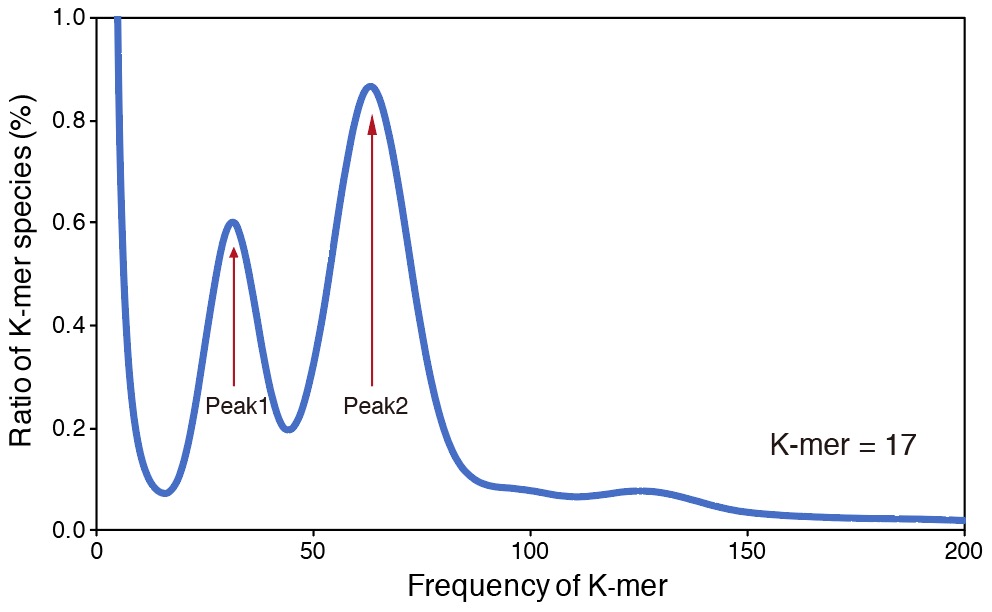

**Supplementary Figure S1**. Distribution of K-mer frequencies in sequencing reads. K-size equal 17.

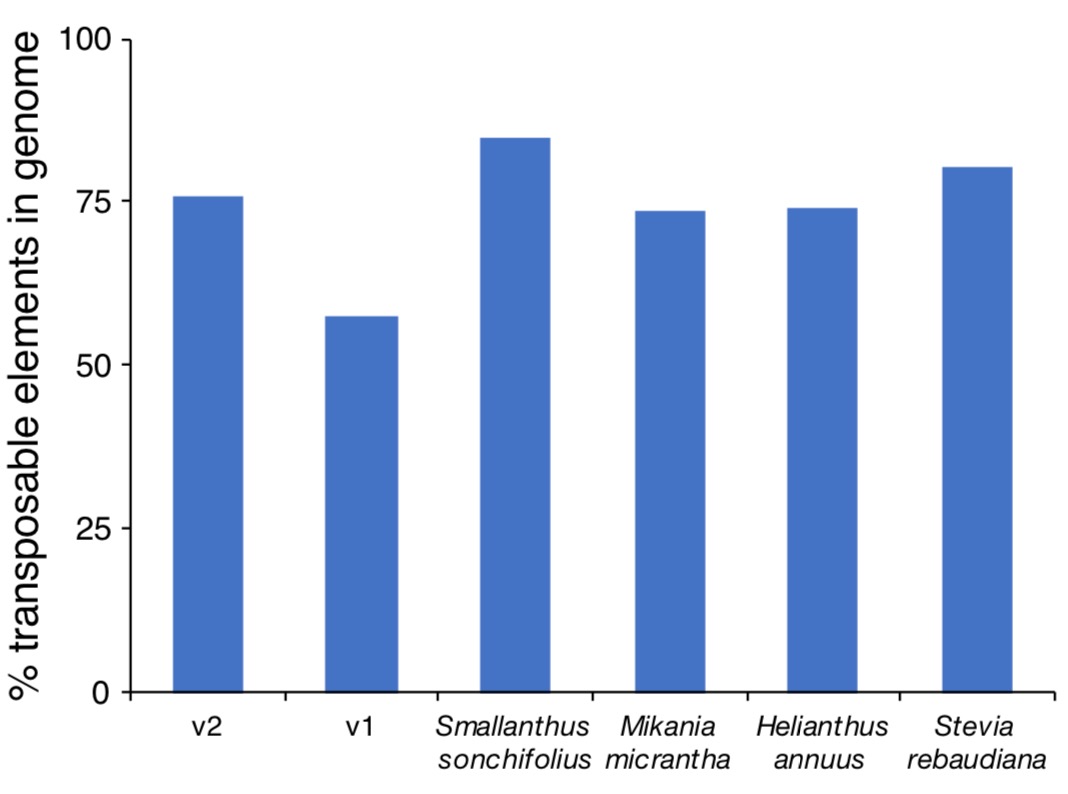

**Supplementary Figure S2.** Comparison of transposable elements among marigold and the closely related species.

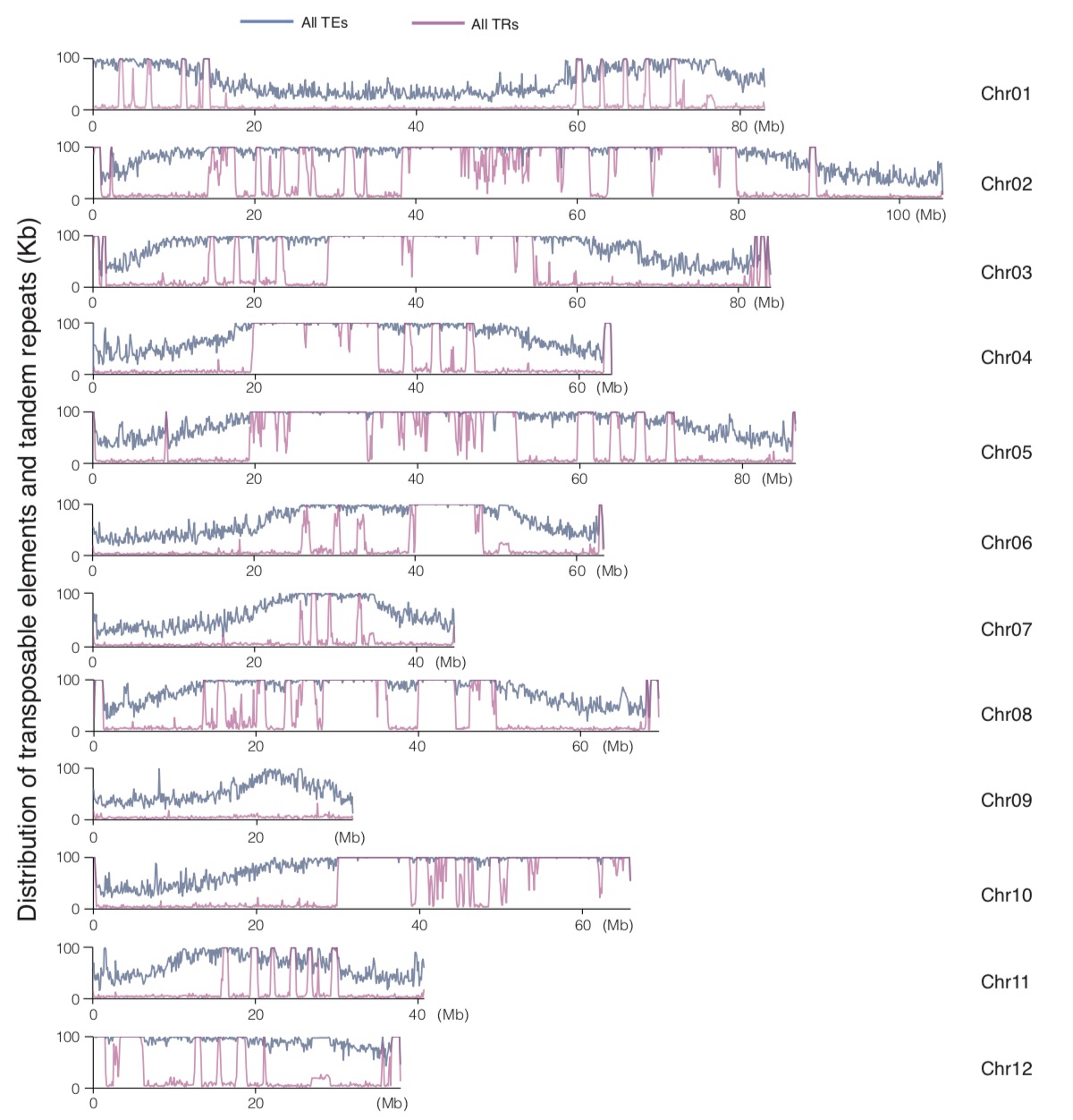

**Supplementary Figure S3**. Distribution of all transposable elements and tandem repeats. We scanned the genome by 100-Kb non-overlapping window as a bin to calculate the length of TEs and TRs.

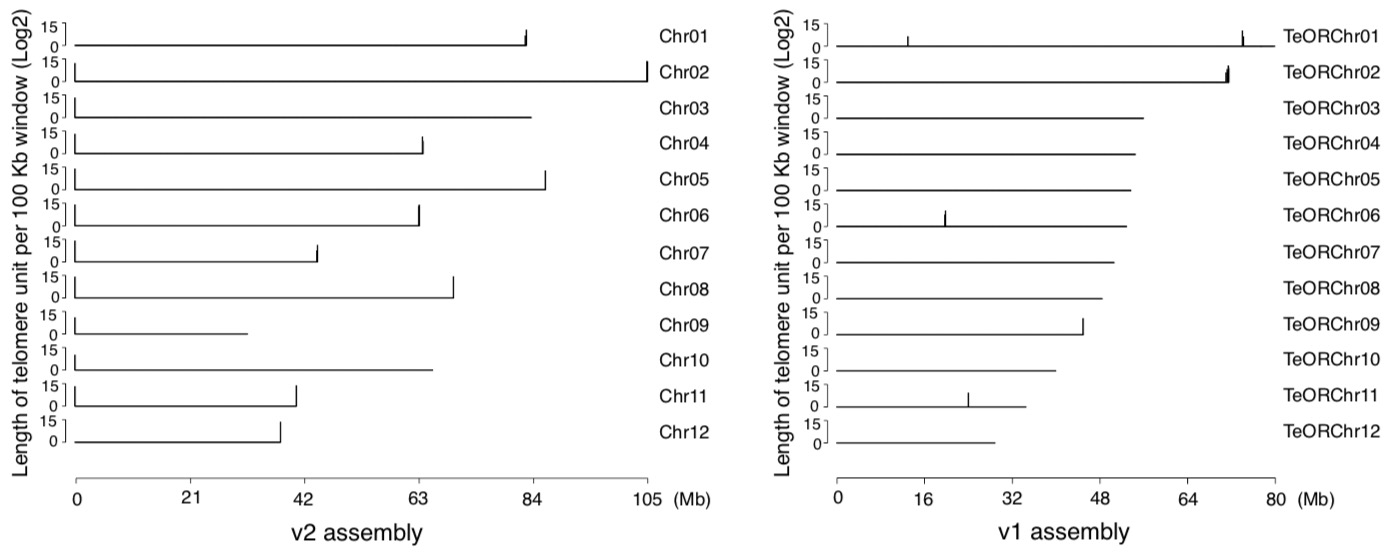

**Supplementary Figure S4**. Distribution of telomeres for the v2 and v1 assembly.

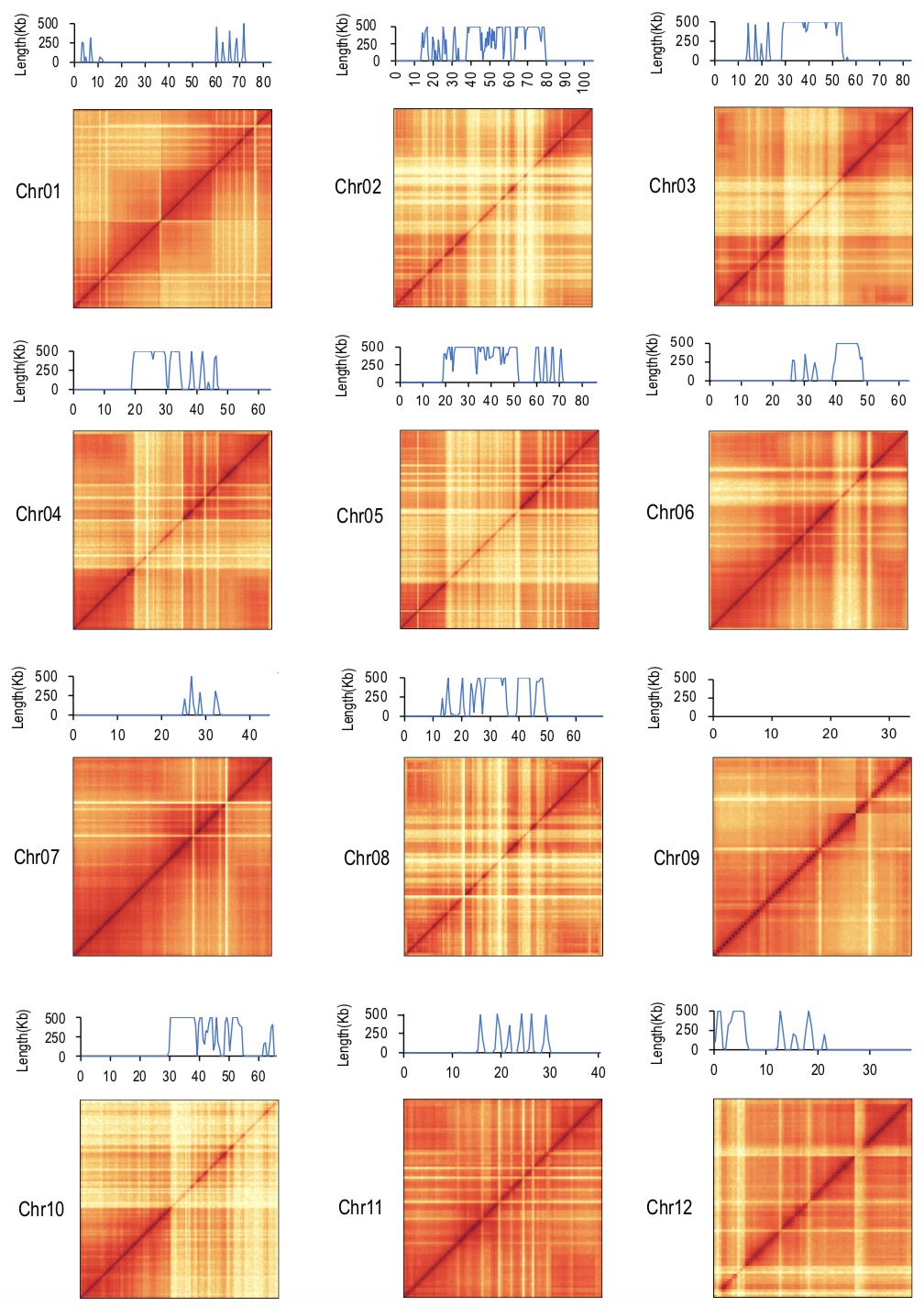

**Supplementary Figure S5.** Distribution of tandem repeats. For each pseudo-chromosome, the above curve showed the distribution of tandem repeats. The window size was 500 Kb. The below showed the heatmap of Hi-C linkage signals. The bin size was 500 Kb.

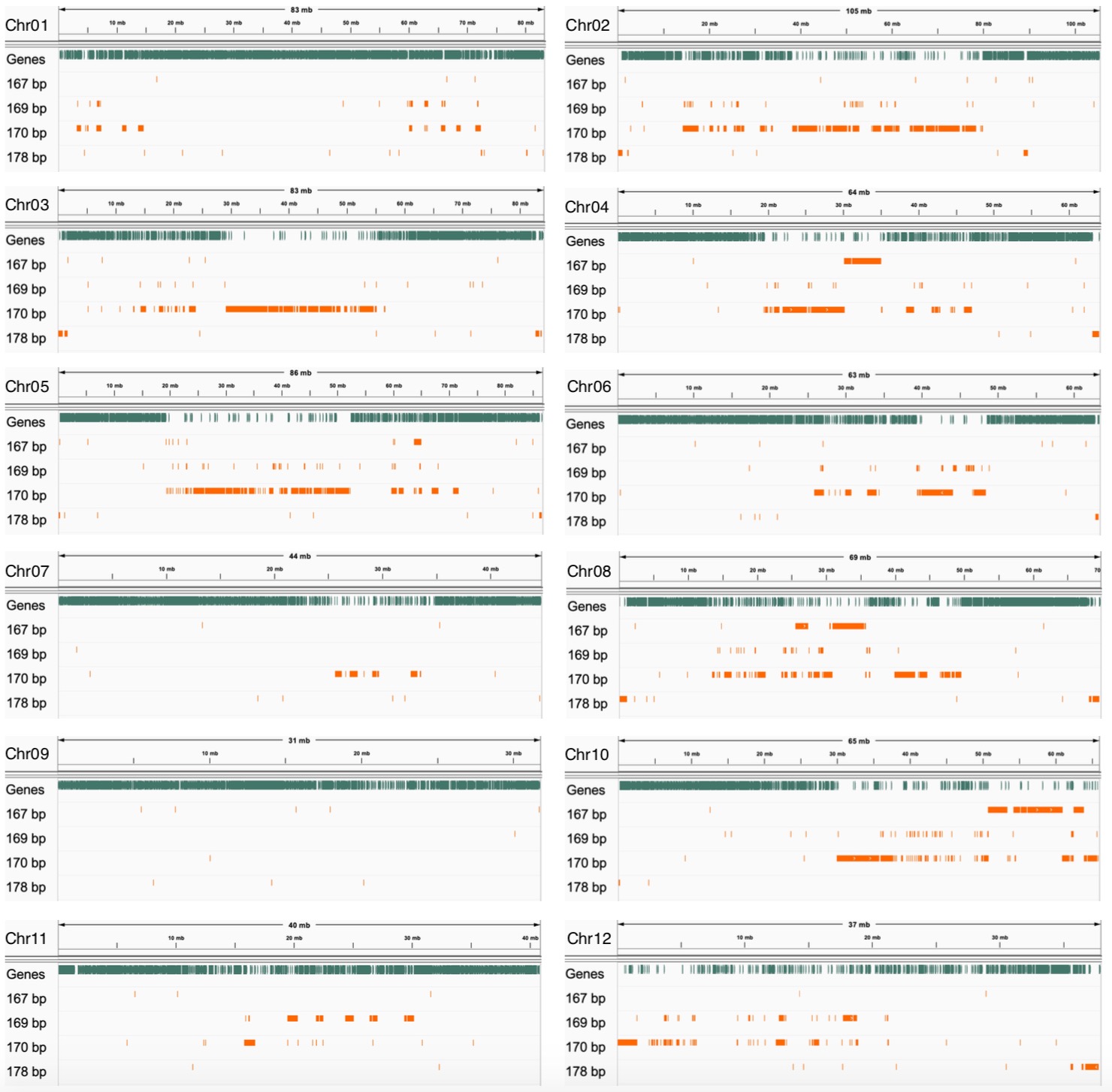

**Supplementary Figure S6.** Views of genes and centromeric repeat units with IGV.

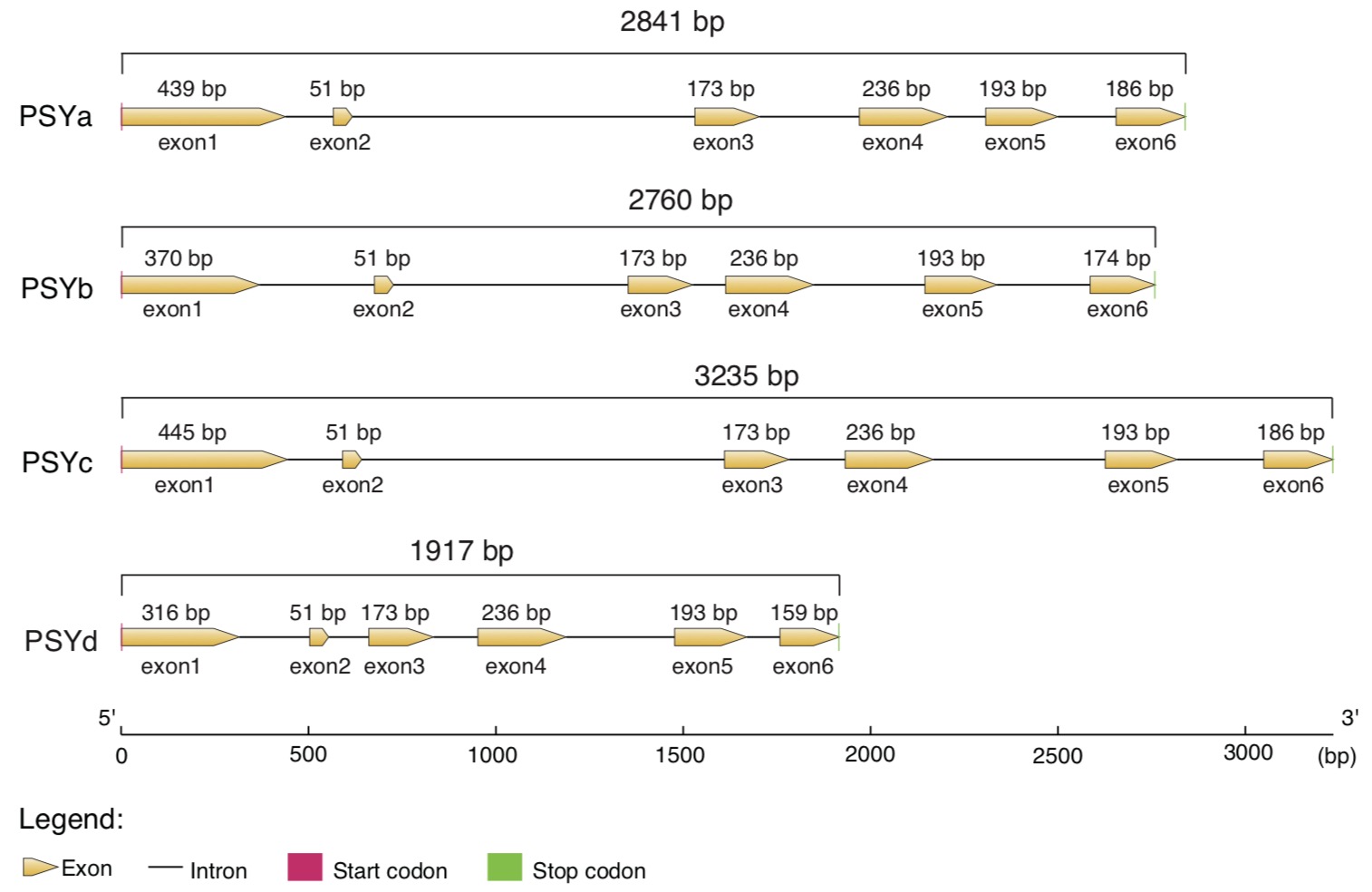

**Supplementary Figure S7.** Comparison of the gene structure for the four *PSY* genes in marigold.

**Supplementary tables**

**Supplementary Table S1.** Summary of the genomic sequencing data for *Tagetes erecta*.

| Type | Sequencing platform | Read number | Base number (bp) | Read N50 length (bp) | Sequencing depth (x) |
| --- | --- | --- | --- | --- | --- |
| PacBio HiFi | PacBio Sequel II | 2,746,197 | 49,699,877,765 | 18,345 | 64 |
| Hi-C | Illumina NovaSeq 6000 | 665,621,308 | 99,843,196,200 | 150 | 128 |

**Supplementary Table S2.** Statistics of primary assembled and purged contigs.

| Statistics | Hifiasm | Purge_dups |
| --- | --- | --- |
| Total contig number | 381 | 47 |
| Total contig length (bp) | 1,010,662,654 | 778,905,021 |
| Maximum length (bp) | 77,349,512 | 77,349,512 |
| Minimum length (bp) | 16,786 | 27,955 |
| N10 (bp) | 65,855,203 | 65,855,203 |
| N20 (bp) | 65,594,574 | 65,594,574 |
| N30 (bp) | 48,669,953 | 51,566,337 |
| N40 (bp) | 43,651,492 | 43,651,492 |
| N50 (bp) | 36,541,802 | 37,953,657 |
| N60 (bp) | 27,378,229 | 27,999,442 |
| N70 (bp) | 17,377,261 | 23,769,366 |
| N80 (bp) | 11,747,071 | 17,377,261 |
| N90 (bp) | 8,775,309 | 10,970,347 |

**Supplementary Table S3.** Statistics of Hi-C data mapping to contigs.

| HiC-pro results | Reads number | Percentage (%) |
| --- | --- | --- |
| Total_pairs_processed | 332,810,654 | 100.00 |
| Unmapped_pairs | 10,650,095 | 3.20 |
| Low_qual_pairs | 173,129,723 | 52.02 |
| Pairs_with_singleton | 57,617,603 | 17.31 |
| Unique_paired_alignments | 91,413,233 | 27.47 |
| Valid_interaction_pairs | 77,585,924 | 23.31 |
| Dangling_end_pairs | 12,120,555 | 3.64 |
| Religation_pairs | 836,062 | 0.25 |
| Self_Cycle_pairs | 84,580 | 0.03 |
| Filtered_pairs | 785,191 | 0.24 |
| Dumped_pairs | 921 | 0.00 |
| Valid_interaction_rmdup | 63,188,228 | 18.99 |

Note: The statistics numbers are obtained from Hi-C pro result files: *.mpairstat, *.mRSstat, and * allValidPairs.mergestat. The “valid interaction rmdup” represents non-redundant and valid Hi-C read pairs, which were used by EndHiC for scaffolding.

**Supplementary Table S4.** Comparison of transposable elements between v2 and v1 assembly.

|  | v2 | | v1 | |
| --- | --- | --- | --- | --- |
| TEs | **Length (bp)** | **Percent (%)** | **Length (bp)** | **Percent (%)** |
| LTR | 500,660,947 | 64.28 | 232,731,398 | 32.91 |
| DNA | 78,601,428 | 10.09 | 19,271,809 | 2.73 |
| LINE | 8,766,338 | 1.13 | 15,870,134 | 2.24 |
| MITE | 383 | 0.00 | ND | ND |
| SINE | 1,319,265 | 0.17 | 0 | 0.00 |
| Others | 1,043,038 | 0.13 | 140,088,732 | 19.81 |
| Total | 590,391,399 | 75.80 | 407,962,073 | 57.69 |

**Supplementary Table S5.** Comparison of predicted genes among marigold and the closely related species.

| Statistics of genes | v2 | v1 | *Smallanthus sonchifolius* | *Mikania micrantha* | *Helianthus annuus* | *Stevia rebaudiana* |
| --- | --- | --- | --- | --- | --- | --- |
| Genome size (Gb) | 0.78 | 0.71 | 2.72 | 1.79 | 3.00 | 1.41 |
| Number of genes | 42,529 | 33,824 | 89,959 | 46,351 | 57,126 | 44,143 |
| Total CDS length (Mb) | 49.27 | 38.80 | 98.19 | 57.59 | 71.30 | 53.58 |
| Percent of total CDS to genome (%) | 6.3 | 5.5 | 3.6 | 3.2 | 2.4 | 3.8 |
| Average CDS length per gene (bp) | 1,158 | 1,147 | 1,091 | 1,242 | 1,248 | 1,213 |
| Average exon number per gene | 5.1 | 5.0 | 5.1 | 5.0 | 4.3 | 5.0 |
| BUSCO assessment (%) |  |  |  |  |  |  |
| Complete | 98.3 | 89.3 | 99.2 | 88.6 | 98.5 | 95.6 |
| Complete and single-copy | 91.4 | 83 | 19.1 | 64.9 | 87.2 | 71.9 |
| Complete and duplication | 6.9 | 6.3 | 80.1 | 23.7 | 11.3 | 23.7 |
| Fragmented | 0.6 | 5.5 | 0.4 | 3.8 | 0.2 | 2 |
| Missing | 1.1 | 5.2 | 0.4 | 7.6 | 1.3 | 2.4 |

**Supplementary Table S6.** Statistics of functional annotation of protein-coding genes.

| Function | Number of genes | Percent of genes |
| --- | --- | --- |
| Protein-coding genes | 42,529 | 100.0 |
| Genes with NCBI-NR hits | 34,159 | 80.3 |
| Genes with KEGG hits | 27,028 | 63.6 |
| Genes with InterPro hits | 36,273 | 85.3 |
| Genes with InterPro terms | 30,489 | 71.7 |
| Genes with GO terms | 22,722 | 53.4 |
| Genes with function terms | 36,877 | 86.7 |

Note: Genes with function terms refer to the genes with at least one term in NCBI-NR, KEGG, InterPro, or GO database.

**Supplementary Table S7.** Statistics of predicted non-coding RNA genes in the genomes of *Tagetes erecta*

| ncRNA genes | Number |
| --- | --- |
| 5S rRNA genes | 527 |
| 5.8S rRNA genes | 493 |
| 18S rRNA genes | 556 |
| 28S rRNA genes | 550 |
| tRNA genes | 1,098 |
| snoR71 | 209 |
| Intron_gpII | 133 |
| U5 | 39 |
| U2 | 38 |
| U1 | 38 |
| U6 | 29 |
| Plant_U3 | 15 |
| MIR169_2 | 15 |
| MIR169_5 | 14 |
| MIR159 | 14 |
| U4 | 13 |
| Plant_SRP | 13 |
| Protozoa_SRP | 12 |
| MIR171_1 | 12 |
| Metazoa_SRP | 12 |
| U3 | 11 |
| mir-395 | 10 |
| Others | 302 |

**Supplementary Table S8.** Statistics of contig and genes for all 12 chromosomes.

|  | v2 | | | | v1 | | | |
| --- | --- | --- | --- | --- | --- | --- | --- | --- |
| Chr_ID | Chr length (bp) | Contig number | Gene number | % genes in Genome | Chr length (bp) | Contig number | Gene number | % genes in Genome |
| Chr01 | 83,052,836 | 4 | 7,725 | 18.16 | 80,606,934 | 122 | 6,459 | 19.10 |
| Chr02 | 105,306,377 | 3 | 3,459 | 8.13 | 72,190,372 | 409 | 2,636 | 7.79 |
| Chr03 | 83,952,692 | 3 | 3,426 | 8.06 | 56,457,020 | 160 | 2,594 | 7.67 |
| Chr04 | 63,998,424 | 5 | 3,216 | 7.56 | 54,910,593 | 164 | 2,667 | 7.88 |
| Chr05 | 86,588,688 | 3 | 3,482 | 8.19 | 54,151,473 | 162 | 2,667 | 7.88 |
| Chr06 | 63,313,408 | 2 | 4,434 | 10.43 | 53,302,000 | 133 | 3,502 | 10.35 |
| Chr07 | 44,643,667 | 4 | 3,803 | 8.94 | 51,000,590 | 76 | 3,318 | 9.81 |
| Chr08 | 69,672,756 | 2 | 2,873 | 6.76 | 48,841,293 | 182 | 2,217 | 6.55 |
| Chr09 | 31,728,754 | 2 | 3,175 | 7.47 | 45,393,979 | 66 | 2,786 | 8.24 |
| Chr10 | 65,855,203 | 1 | 2,701 | 6.35 | 40,397,928 | 191 | 2,028 | 6.00 |
| Chr11 | 40,789,275 | 2 | 3,065 | 7.21 | 34,835,773 | 45 | 2,483 | 7.34 |
| Chr12 | 37,869,363 | 2 | 866 | 2.04 | 29,107,531 | 102 | 467 | 1.38 |
| Total | 776,771,443 | 33 | 42,225 | 99.29 | 621,195,486 | 1,812 | 33,824 | 100.00 |

**Supplementary Table S9.** Statistics of 4 centromeric repeat units.

| Repeat units | Copies | Total length | % genome |
| --- | --- | --- | --- |
| 170-bp | 682,138 | 115,963,477 | 14.89 |
| 167-bp | 127,857 | 21,352,186 | 2.74 |
| 178-bp | 55,092 | 9,806,340 | 1.26 |
| 169-bp | 40,680 | 6,874,852 | 0.88 |

**Supplementary Table S10.** Statistics of the centromeric repeat units for all 12 pseudo-chromosomes.

|  | 167-bp unit | | 169-bp unit | | 170-bp unit | | 178-bp unit | |
| --- | --- | --- | --- | --- | --- | --- | --- | --- |
| Chr | Copies | Length (Kb) | Copies | Length (Kb) | Copies | Length (Kb) | Copies | Length (Kb) |
| Chr_01 | 8 | 1 | 2,269 | 383 | 22,726 | 3,857 | 90 | 16 |
| Chr_02 | 26 | 4 | 1,553 | 259 | 158,015 | 26,853 | 10,454 | 1,861 |
| Chr_03 | 11 | 2 | 112 | 19 | 122,244 | 20,803 | 10,581 | 1,885 |
| Chr_04 | 25,716 | 4,297 | 358 | 61 | 60,244 | 10,162 | 4,821 | 862 |
| Chr_05 | 7,844 | 1,325 | 610 | 103 | 116,879 | 19,869 | 3,320 | 592 |
| Chr_06 | 14 | 2 | 6,933 | 1,171 | 39,125 | 6,645 | 2,136 | 381 |
| Chr_07 | 5 | 1 | 2 | 0 | 9,331 | 1,584 | 105 | 19 |
| Chr_08 | 34,558 | 5,793 | 3,310 | 561 | 58,603 | 9,957 | 12,447 | 2,220 |
| Chr_09 | 13 | 2 | 2 | 0 | 2 | 0 | 7 | 1 |
| Chr_10 | 59,651 | 9,986 | 2,447 | 416 | 72,276 | 11,892 | 1,365 | 243 |
| Chr_11 | 7 | 1 | 15,676 | 2,655 | 3,924 | 665 | 4 | 1 |
| Chr_12 | 5 | 1 | 7,408 | 1,256 | 18,770 | 3,070 | 7,338 | 1,308 |
| Total | 127,857 | 21,416 | 40,680 | 6,884 | 682,138 | 115,357 | 52,668 | 9,389 |
